# The structural basis of long-term potentiation in hippocampal synapses, revealed by EM-imaging of lanthanum-induced synaptic vesicle recycling

**DOI:** 10.1101/2022.04.22.489201

**Authors:** John E. Heuser

## Abstract

Hippocampal neurons in tissue-culture were exposed to the trivalent cation lanthanum for short periods (15 to 30 minutes) and prepared for electron microscopy (EM), to evaluate the stimulatory effects of this cation on synaptic ultrastructure. Not only were characteristic ultrastructural changes of exaggerated synaptic vesicle turnover seen within the presynapses of these cultures - - including synaptic vesicle depletion and proliferation of vesicle-recycling structures - - but the overall architecture of a large proportion of the synapses in the cultures was dramatically altered, due to large postsynaptic ‘bulges’ or herniations into the presynapses. Moreover, in most cases these postsynaptic herniations or protrusions produced by lanthanum were seen by EM to distort or break or ‘perforate’ the so-called postsynaptic densities (PSD’s) that harbor receptors and recognition-molecules essential for synaptic function. These dramatic EM-observations lead us to postulate that such PSD-breakages or ‘perforations’ could very possibly create essential substrates or ‘tags’ for synaptic growth, simply by creating fragmented free-edges around the PSD’s, into which new receptors and recognition-molecules could be recruited more easily, and thus they could represent the physical substrate for the important synaptic-growth process known as “long-term potentiation” (LTP). All of this was created simply in hippocampal tissue-cultures, and simply by pushing synaptic vesicle recycling way beyond its normal limits with the trivalent cation lanthanum; but we argue in this report that such fundamental changes in synaptic architecture - - given that they can occur at all - - could also occur at the extremes of normal neuronal activity, which are presumed to lead to learning and memory.

## INTRODUCTION

Of all the important structural features of the synapse that have been viewed in the electron microscope over the decades, the one feature that has been the most correlated with LTP, and especially in the context of EM-preparations generated from hippocampal tissues that were experimentally manipulated into LTP-conditions, that one single feature would have to be the *perforated postsynaptic density* (or *perforated PSD*).

It was the life’s work of one of the leading electron microscopists of the hippocampus, Yuri Geinesman, to establish this correlation (**Reference Set #1 (Geinesman).** Another top electron microscopist of the hippocampus, Kirsten Harris, reported *perforated PSD’s* almost as often, but remained more agnostic about their role in hippocampal LTP (**Reference Set #2 (Harris).** Likewise, two other top EM-labs that published a lot on the hippocampus, Michael Stewart’s in Milton Keynes, England, and Dominique Muller’s in Geneva, Switzerland, reported finding *perforated PSDs* in many, many publications and also made outstanding contributions toward understanding how these might be involved in LTP or ‘synaptic learning’ (**Reference Set #3 (Stewart); Reference Set #4 (Muller)**. All this history culminated with the later, outstandingly beautiful cryo-EM work on this same topic, which came from Michael Frotscher’s lab in Freiberg, Germany (**Reference Set #5 (Frotscher).** Along the way, and right up to the present day, many other EM-labs have weighed in with valuable correlative data and their own ideas on how LTP might come about (**Reference Set #6 (Other EM-studies of hippo learning),** and numerous reviews of perforated synapses have been published (although without any serious attempt to correlate them with learning or memory) **(Reference Set #7 (Petralia&Yao reviews).**

At hippocampal synapses, the postsynaptic densities or PSD’s are ordinarily disk-shaped entities that are generally located directly across from presynaptic neurotransmitter release-sites **Reference Set #8 (PSD basic structure).** They generally appear almost *continuous* in thickness and density across the breadth of the disk, and even though recent close inspections have begun to suggest that each PSD-disk may have a sub-structure, and may be composed of sub-modules or ‘nanomodules’ **Reference Set #9 (synaptic ‘modules’ via LM-viewing**), these PSD-components pack closely enough together to create the general impression in the electron microscope (EM) of a continuous plaque or disk.

‘Perforated’ PSDs, in contrast, are variable in outline. They look more like irregular and discontinuous patches in the EM, but they are presumed to be composed of the same components that are found in the more normal, continuous PSD-disks. Reviewing the aforementioned references, one can find a wide range of thoughts about them, everything from conclusions that such ‘perforated’ PSDs are in the process of division of one plaque into two (ultimately to form two different postsynaptic spines, in some of the bolder claims), to conclusions that they are simply a by-product of synaptic activity. But overall, ‘perforated’ PSDs have generally been interpreted as synapses-in-augmentation, as would be expected if they were the structural correlates of the synaptic enhancement that is presumed by everyone to be the fundamental basis of LTP **Reference Set #10 (Background of Long Term Potentiation).**

In this report, we present serendipitous observations that could help to explain exactly <u>how</u> “bursts” of presynaptic activity could cause perforated PSDs to form, in the first place. Specifically, we argue that these bursts of presynaptic activity could create delays in the process of synaptic vesicle recycling, which could make the presynapse expand in surface area, and do in such a manner that it could actually strain and break the normal plaque-like PSD into a ‘perforated’ PSD.

Our EM-images were obtained from primary (dissociated-cell) hippocampal cultures prepared by classical techniques **Reference Set #11 (Methods for hippocampal slices & cultures).** These were chemically stimulated *via* direct application of low doses (0.1mM) of the trivalent cation, lanthanum (La+++). This magical trivalent cation has been used to stimulate spontaneous neurosecretion in many different preparations over the past half century **Reference Set #12 (La++ stimulates all sorts of secretion),** but still, no one knows exactly *how or why* it does so, especially because it is generally considered to be a “super-calcium” that actually *blocks* most calcium-channels, and thus blocks most calcium-evoked neurosecretory phenomena, it doesn’t *stimulate* them **Reference Set #13 (La+++ blocks Ca++ channels).**

When we first faced this conundrum fifty years ago (the conundrum of <u>why</u> lanthanum blocks calcium-induced neurosecretion but hugely stimulates *spontaneous* transmitter release, in the form of huge bursts of miniature endplate potentials or “m.e.p.p.’s” at the NMJ), we simply could not explain it, but nevertheless we ‘used’ the phenomenon to deplete synaptic vesicles from frog neuromuscular junctions, and thus to expose for the first time the resultant membranous-transformations that turned out to represent synaptic vesicle recycling **Reference Set #14 (Heuser&Miledi - discovery of SV recycling via La+++ stim).** Our scientific colleagues in those days quickly followed suit, and <u>also</u> used lanthanum to show that a dramatic expansion of the presynaptic membrane can accompany this spontaneous lanthanum-induced transmitter discharge, due to a slow-down of synaptic vesicle recycling and accumulaton of vesicle membrane on the presynaptic surface **Reference Set #15 (La+++ moves SV membrane to presynaptic surface).** In fact, we intend to show here -- by classical thin-section EM -- that it is this presynaptic *expansion* which is most dramatic in hippocampal cultures, and furthermore, that the unique distortion(s) which this expansion creates on the postsynaptic side of hippocampal synapses can greatly help to explain *how and why* their PSD’s can become perforated during enhanced synaptic activity.

## RESULTS

*(See all Figures and their legends, for the demonstrations of the points made here.)*

**Figure 1.**
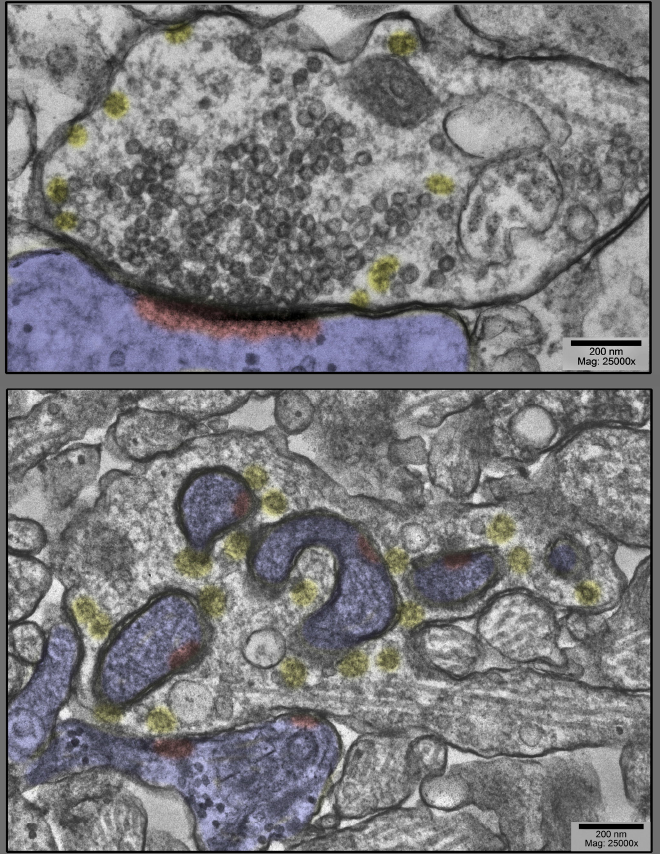
<u>Upper panel</u> shows a prototypical spine-synapse from a control hippocampal culture, a culture not stimulated at all. Abundant synaptic vesicles are collected on the presynaptic side, converging on a solid, disk- or plaque-like postsynaptic density (highlighted red, inside the postsynapse, which is highlighted blue in this and all subsequent figures). Only a few clathrin-coated vesicles are present (highlighted yellow), and only at the periphery of the vesicle cluster. <u>Lower panel</u> shows, by way of contrast, a dramatically different spine-synapse from a hippocampal culture exposed to 0.1mM La+++ for 20 min @ 37°C. Synaptic vesicles are severely depleted, clathrin-coated vesicles (yellow) are unusually abundant, and the postsynapse (highlighted blue) has ‘protruded’ into the presynapse in several places, dragging along only fragments of the PSD (highlighted red).

**Figure 2.**
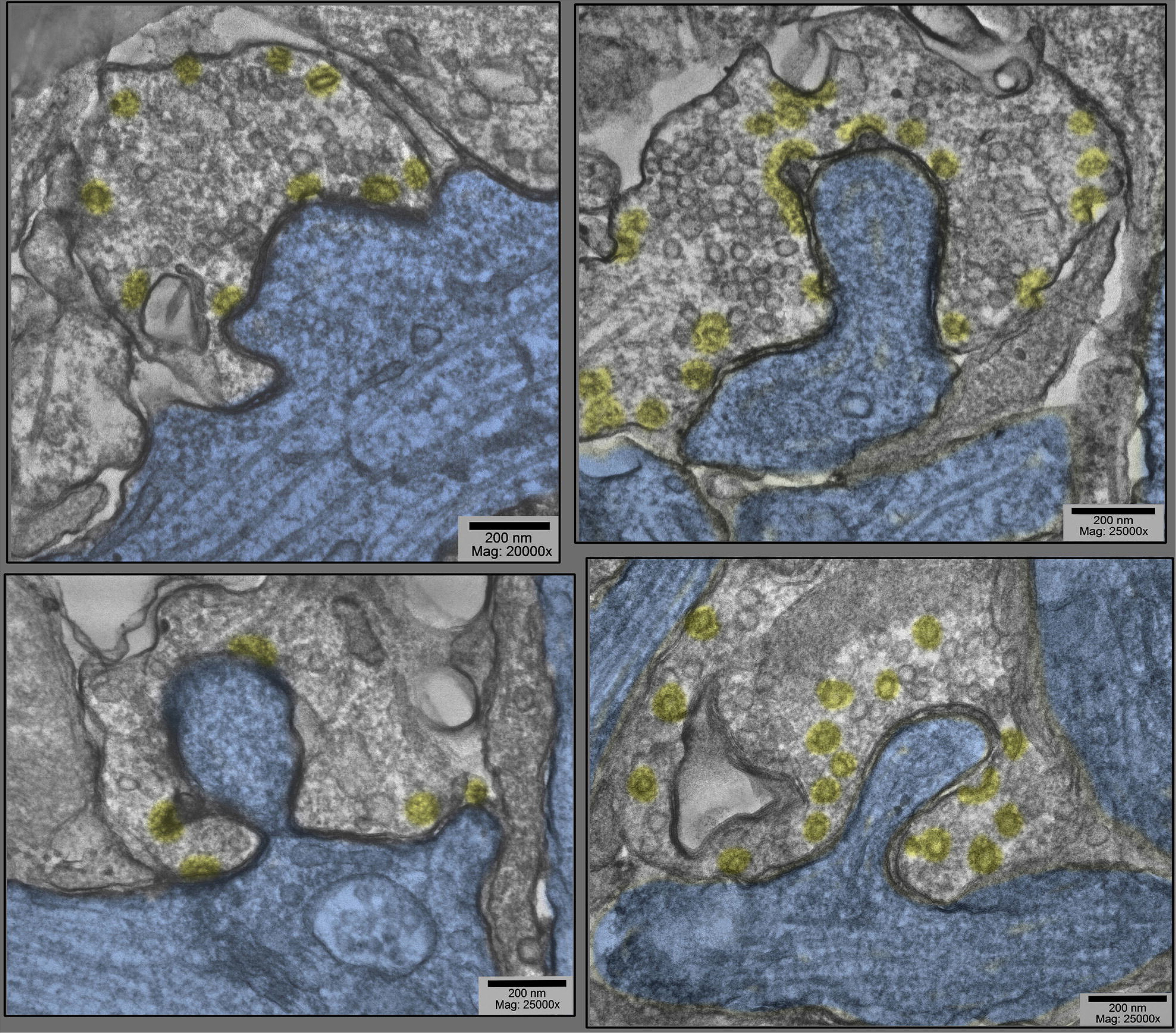
Examples of dendritic-shaft synapses in hippocampal cultures exposed to 0.1mM La+++ for 10-20 min @ 37°C. Synaptic vesicles are more or less depleted; but in all cases, clathrin coated vesicles (presumably involved in the recycling of synaptic vesicle membrane) are increased in abundance, especially along the borders of the postsynaptic ‘protrusions’ (highlighted blue). Being shaft-synapses (thus presumably inhibitory or Grey’s type II), the PSD’s are not prominent in these fields and thus are not highlighted red.

**Figure 3.**
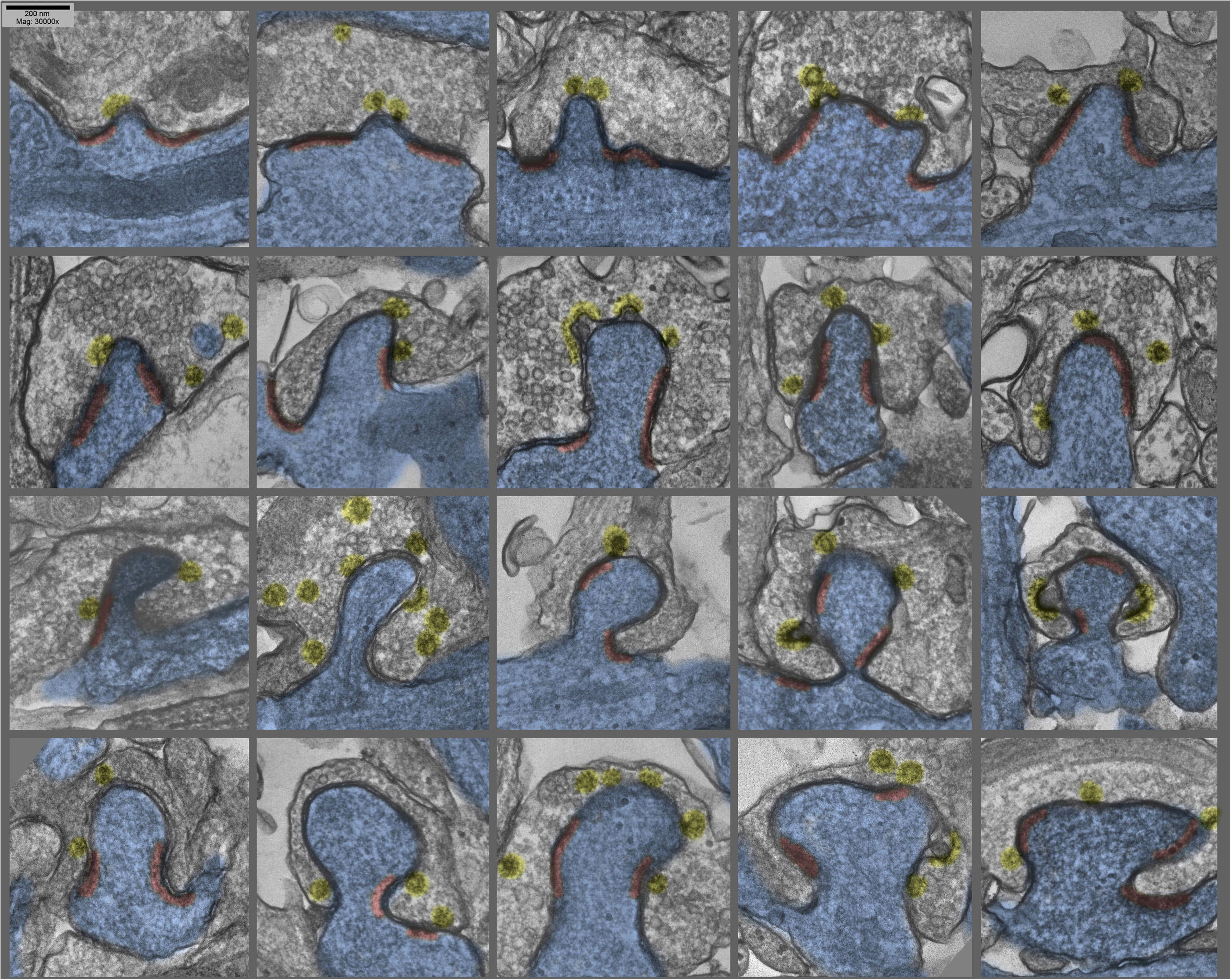
Montage of several different postsynaptic ‘protrusions’ (or presynaptic ‘hugs’, depending on one’s vantage-point) in hippocampal cultures exposed to 0.1mM La+++ for 10-20 min @ 37°C. Synaptic vesicles are more or less depleted, and clathrin coated vesicles (again highlighted yellow) are relatively abundant, and PSD’s (highlighted red) are hanging back or are fragmented and ‘holding on’ to the edges of the postsynaptic protrusions.

**Figure 4.**
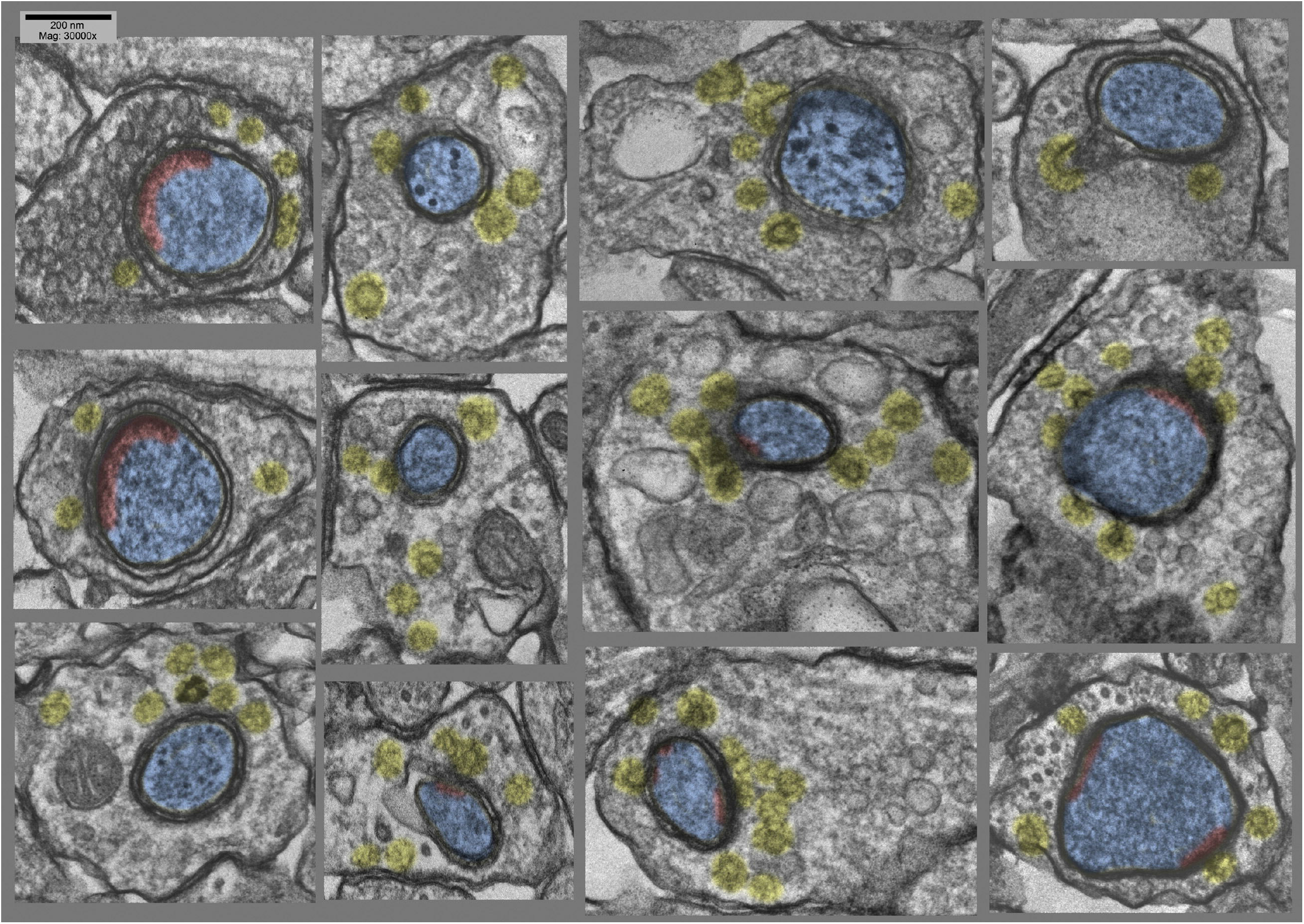
Montage of <u>cross-sections</u> of several other postsynaptic ‘protrusions’ or presynaptic ‘hugs’, from the same hippocampal cultures as in Fig. 3 (exposed to 0.1mM La+++ for 10-20 min @ 37°C). Again, it is quite apparent that synaptic vesicles are more or less depleted, compared to the relative abundance of clathrin-coated vesicles (highlighted yellow). Again, PSD’s (highlighted red) appear to be relatively fragmented or absent from these deeper regions of the postsynaptic involutions.

**Figure 5.**
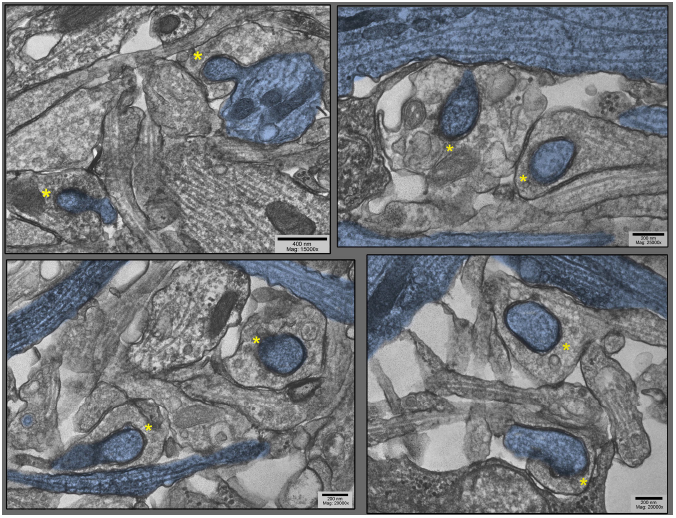
Lower-magnification survey-views of hippocampal cultures exposed to 0.1mM La+++ for 10-20 min @ 37°C), where only the particular dendrites that are involuting into their apposed synapses are highlighted blue (other dendrites are present in these fields, as well). These views were chosen specifically to demonstrate the old adage that George Palade, the great ‘founder’ of biological electron microscopy, always stressed: namely, that if a structural feature could be found *two or more times in one and the same field-of view* in the electron microscope, then it must be generally present, and must be not an artifact. <u>Two</u> postsynaptic ‘protrusions’ are indicated by asterisks in each field.

**Figure 6.**
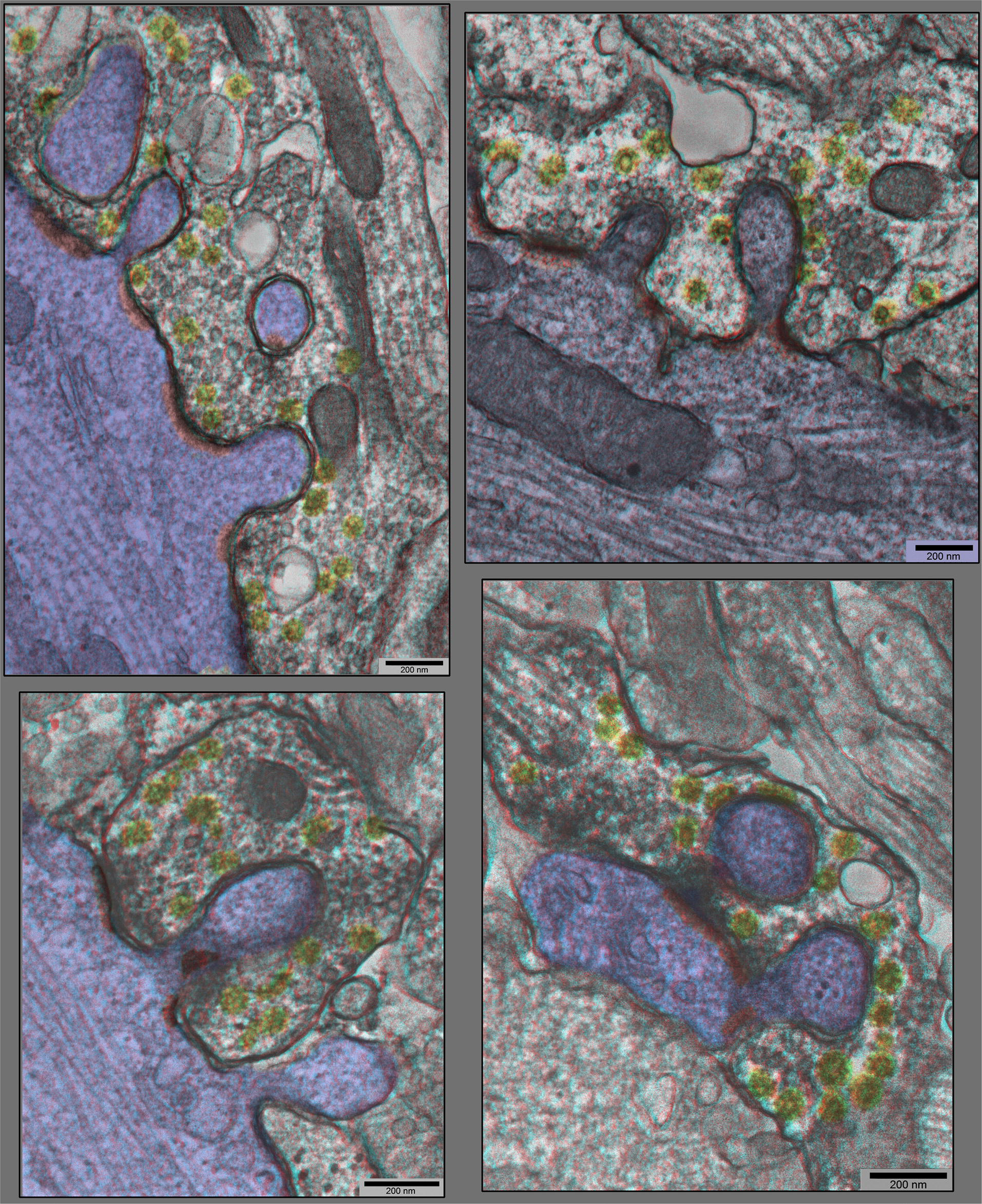
”Anaglyph” 3-D views of additional synapses in hippocampal cultures exposed to 0.1mM La+++ for 10-20 min @ 37°C), from plastic blocks that were cut thicker (at 120-150nm), and photographed at +20° & −20° of tilt in the EM, then superimposed to make the 3-D ‘anaglyphs’. These require red/green ‘anaglyph’ glasses to fully appreciate, but even without, it is still readily apparent that synaptic vesicles are relatively depleted, coated vesicles are relatively abundant (yellow), and postsynaptic densities are relatively fragmented (PSD’s red, on dendrites highlighted blue).

**Figure 7.**
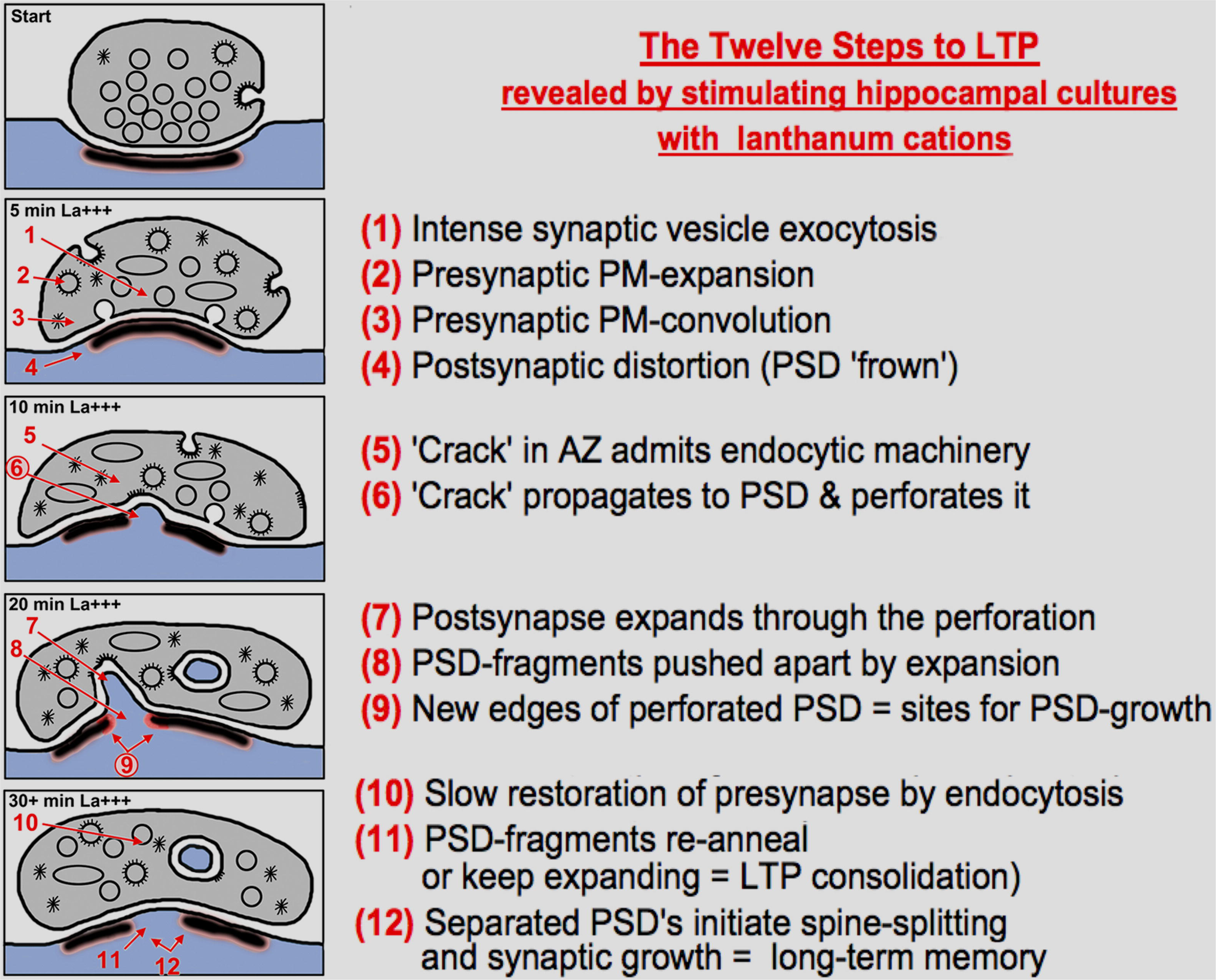
Diagrammatic summary of the observations, interpretations, and hypotheses presented in this study, presented as ‘twelve steps,’ to paraphrase a term from popular culture. The diagram is self-explanatory, but many steps will need to be validated or explained mechanistically, by future work…especially the idea that enhanced endocytosis can cause a ‘break’ in the presynaptic active zone (AZ), and that this break can propagate to the PSD to begin the process of perforation or intrusion or spinule-formation.

### 1. Structural evidence of lanthanum’s stimulatory-effects

Fifty-seven separate experiments were performed with lanthanum on hippocampal cultures, adjusting the time of exposure (5-30 min), the dose of lanthanum (0.1mM-1mM), the level of calcium-counter ions (zero to 1mM), and the method of fixation for EM (glutaraldehyde in cacodylate buffer *vs.* in Hepes buffer). Such classical dissociated-cell hippocampal cultures invariably display a huge range of synaptic types, only vaguely reflecting the characteristic and stereotypical forms of synapses observed in the intact hippocampus or in hippocampal slices **Reference Set #11 (Methods for hippocampal slices & cultures, OP CIT**). Nevertheless, most of the synapses that manage to reform in such dissociated-cell cultures fall into two major categories: <u>bouton-like</u> (e.g., onto dendritic protrusions that are generally described as primitive ‘spines’), and <u>plague-like</u> (e.g., directly onto the shafts of the primitive, disoriented dendrites in such cultures). The former types of synapses typically have single PSD plagues that are generally considered to be ‘excitatory.’ These are the so-called ‘*asymmetric*’ synapses, due to their relatively thick PSDs. The latter typically display 2 or 3 adjacent plaques in the dendritic shaft and are generally considered to be ‘inhibitory’ These are the so-called ‘*symmetric*’ synapses, due to their relatively thin and often barely perceptible PSDs (**Reference Set #8 (PSD basic structure) OP CIT**).

Lanthanum promptly changes the appearance of both bouton-like and plaque-like presynaptic terminals, replacing many of their synaptic vesicles with clathrin coated vesicles, even in the first 5-10 min after application, and at later times, leaves many of these presynaptic terminals almost empty, or at best, partially filled with irregular membrane forms (plus residual clathrin coated vesicles and ‘empty cages’). Essentially, these are the structural changes that we observed originally at the frog NMJ (**Reference Set #14 (Heuser&Miledi - SV recycling in La+++) OP CIT)**, which initiated the whole idea that synaptic vesicle membrane could be recycled…structural changes that we have recently been able to show occur in mammalian NMJs, as well (**Reference Set #16 (Heuser & Tenkova intro of ear-muscle NMJs).**

### 2. Consequences of lanthanum’s stimulatory-effects projected onto the postsynapse

Most unexpected, however, was the dramatic change in the <u>overall configuration</u> of these cultured synapses, a configuration that began to appear in lanthanum at 10 min, and peaked in abundance at 20 min, (and seemed to die down by 30 min). This was manifest as a discrete ‘bulge’ of the postsynapse, directly into the midst of the presynaptic terminal.

Generally, the postsynaptic protrusions observed in lanthanum occupy a considerable portion of the presynaptic cytoplasm, and draw a considerable mass of postsynaptic cytoplasm into them. In the EM, this bolus of cytoplasm generally appears featureless, or appears finely mesh-like in appearance. It does not look like an active, actin-based <u>growth</u> from the postsynapse, but more like a passive inclusion, only occasionally containing any recognizable postsynaptic membranous organelle.

It is important to consider one clue as to *how or why* these protrusions form in the first place. This comes from closely examining the membrane of the *presynapse* that forms the protrusion (or we could say in the *invagination*, when considered from the presynaptic side). It typically displays all the ‘spikes’ and ‘clathrin-cage-fragments’ seen in other regions of the presynapse that are undergoing rapid and abundant clathrin-coated vesicle formation. That is, it looks in the EM as if the presynaptic membrane is “committed to endocytosis” in these involutions, or is “trying” to do endocytosis in the regions that have been drawn inwards into the protrusion.

### 3. Unique positioning of the postsynaptic protrusions and distortions

Perhaps the most important aspect of these postsynaptic protrusions in the present context, however - - that of considering their possible role in LTP - - is that they typically occur *right in the midst* of the PSD. As a consequence, they occasionally drag portions of the PSD inward as they form, depositing these PSD fragments along the ‘necks’ of the invaginations. More often, however, PSD components appear to be <u>excluded</u> from these invaginations - to somehow “hang back” - such that the PSD becomes <u>perforated or partitioned</u> by the invagination.

How or why these invaginations form *right in the midst* of the PSD - - rather than around its edges, for example, where one might imagine the membranes to be more ‘flexible’ - - is one of the great mysteries that emerges from this study, and remains to be answered.

## DISCUSSION

### 1. Clues about how and why the postsynaptic protrusions develop

The hypothesis that emerges from these observations is that overly exuberant presynaptic expansion and attempts at endocytosis are ‘deforming’ their active zones and producing the inward membrane-bulges that perforate the PSD. Support for this hypothesis came from our attempts to duplicate the effects of lanthanum on these cultures with other sorts of chemical stimulation of them (either by the applying the excitatory neurotransmitter NMDA, or by elevating potassium to depolarize all the cells in the culture). But <u>neither</u> of these forms of stimulation induced <u>any</u> such protrusions or inward invaginations of their postsynapses, <u>at all</u>. The most they showed was the slight change in PSD-curvature that has been reported before **Reference Set #17 (PSD curvature** Δ**s).** We would argue that this was probably because the synapses in these NMDA or K+ stimulated cultures were not driven out of their “comfort zones,” the zones where they could adequately compensate for their enhanced transmitter release (and their enhanced synaptic vesicle exocytosis) by commensurately accelerating their membrane recycling processes, so they stayed effectively ‘in balance’, and did not develop any net accumulation of synaptic vesicle membrane on their surfaces, and thus did not expand in surface-area (and consequently, did not attempt to “embrace” the postsynapse or support any postsynaptic protrusions).

Indeed, we may learn someday that the most important reason for why lanthanum-stimulation is so effective at expanding the surfaces of nerve terminals and bringing out signs of synaptic vesicle recycling is that it somehow <u>slows down</u> this membrane recycling, at the same time that it stimulates transmitter-release, itself. This may someday be explained by several different mechanisms: for example, by some sort of ‘stiffening’ of the presynaptic membrane or by La+++ tending to ‘glue’ the presynaptic membrane to the extracellular matrix, or possibly by its blocking sodium and calcium-entry through the presynaptic membrane, which may in some way *directly* slow down some aspect of the recycling-process. Indeed, there is a large body of evidence which suggests that clathrin coated vesicle formation and/or synaptic vesicle recycling is somehow dependent on intracellular Ca++ being at just the right level (**Reference Set #18 (Ca++-dependence of synaptic vesicle recycling).** In any case, it is abundantly clear from the present observations and from past work that lanthanum somehow creates a greater imbalance between exocytosis and endocytosis than any other form of synaptic stimulation, and consequently, produces the most enhanced accumulation of synaptic vesicle membrane on the presynaptic surface, and thus, the greatest expansion of the presynaptic membrane.

In this respect, these synaptic perturbations in lanthanum-stimulated hippocampal cultures show a remarkable parallel with the abundant, multiple, and florid invaginations of the presynaptic membrane that were seen so many decades ago in frog NMJ’s treated with lanthanum (**Reference Set #15 (La+++ moves SV membrane to presynaptic surface) OP CIT**). These were *also* studded with endocytic profiles, and the thinking at that time was the same as today - - that lanthanum caused such an unremitting stimulation of the NMJ, that endocytosis became exhausted and ‘blocked’ at that point, or at least greatly slowed down, such that it left much of the discharged synaptic vesicle membrane on the surface of the presynaptic terminal, and thereby *expanding* that surface. A lot of good immunocytochemistry was done in the ensuing decades, which entirely supported this view (**Reference Set #15, OP CIT**).

The important difference between the changes seen here, in lanthanum-stimulated hippocampal synapses, compared to the old NMJ observations, is that <u>here</u> the *postsynapse proper* is pulled into the presynapse. This cannot happen at the NMJ, where the presynapse is separated from the postsynapse by a thick and rigid basal lamina. Instead, at the NMJ the surrounding Schwann cell gets pulled into the invaginations -- or one could say, the Schwann cell ends up ‘protruding’ into the presynaptic terminal. ((In this regard, it is worth noting that we observed no such “drawing-inward” of surrounding glial processes in any of our lanthanum-stimulated hippocampal cultures. This at least partly due to the simple fact that there are not very many glial processes around the synapses in our cultures, in the first place; and the few glial cells that <u>do</u> happen to be there may have very little ‘give’.)) Quite different is the situation at all NMJs, where Schwann cells totally embrace the whole nerve terminal, except at its immediate contact with the muscle, and where the Schwann cells are highly redundant and ‘plastic’, so they actively fill in any convolutions of the presynaptic membrane that develop during stimulation. Interestingly, in the primary neuron cultures from the De Camilli lab, their endocytosis-inhibited genotypes show <u>both</u> glial and postsynaptic involutions when the synapses are stimulated (**Reference Set #15a (De Camilli lab’s endocytosis-retarded neuron-cultures)).**

### 2. Comparing classical postsynaptic “spinules” with the protrusions and distortions seen here in lanthanum

There is no good reason to think that the so-called synaptic “spinules” described in many previous studies of hippocampal synapses (**Reference Sets 1-4, OP CIT**) can or should be differentiated from the *fatter* postsynaptic protrusions into the presynapse that we are describing here, as being the consequences of lanthanum-stimulation. *Neither* type of invagination contains any postsynaptic structure that would suggest they were ‘active’ invasions into the presynapse. In other words, *neither* type contains any signs of actin, nor any other cytoskeletal component that might suggest that they actively <u>push</u> their way into the presynapse. Instead, the EM’s presented here show clearly that both ‘spinules’ and their fatter counterparts invade regions of the presynapse that show all the signs of being engaged in clathrin-mediated endocytosis. As explained above, this dedication to endocytosis is reason enough to understand why the presynapse should be <u>involuted</u> at those sites. But it also suggests that the spinules and their fatter counterparts, the postsynaptic protrusions, are not *actively splitting* the PSDs as they invade the presynapse. Again, their lack of contractile/propulsive machinery would seem to rule this ou. Instead, this PSD-splitting appears to be a *passive* process, almost an inadvertent consequence of the presynaptic involution, itself (inadvertent, except that it perhaps was selected by nature to be the fundamental ‘growth’-event of LTP !!)

It is worth stressing here that both ‘spinules’ and the fatter protrusions display uniformly close approximations of pre- and post-membranes, in their midst - - the important point being that these are *closer* approximations than are observed at the AZ/PSD differentiations of the synapse *per se*, where so many spanning and attachment molecules are known to be located (and are known to be so abundant and strong that they can even hold the pre- and post-together during homogenization of nervous tissue) (**Reference Set #8 (PSD basic structure), OP CIT**). All these specific synaptic attachment-proteins have ∼15-20nm spanning-lengths, which are *greater* than the pre-to-post membrane-separation seen at the protrusions/involutions under question (∼10nm). Indeed, this 10nm separation-distance that we observe in all the protrusion-sites is the normal ‘minimum’ found everywhere in the neuropil of our hippocampal cultures - - e.g., between glia and neuronal elements, between undifferentiated neural elements and each other, etc., etc.. (Of course, this ‘minimum’ is not *nearly* close enough to suggest electrical coupling, or anything of that sort.) In any case, this is one more indication that the protrusions are not pulling in the PSDs and their associated attachment-proteins with them; rather, they are *splitting* the PSDs, and leaving these attachment-proteins behind.

Here, we should add the qualification that despite the obvious structural parallels between synaptic spinules and their fatter counterparts displayed here, nothing in our studies to date would suggest that a precursor/product relationship exists between them. While both seem clearly to be exacerbated or accentuated by synaptic stimulation, the *timing* of this stimulation varies over several orders of magnitude (from just a few minutes in the present report of acute, ongoing chemical stimulation) to hours or longer, in previous studies characterizing the long-term after-effects of LTP-induction, as listed above. On other words, it is not (yet) possible to suggest that spinules grow into fatter protrusions as they invade, or that fat protrusions shrink down to spinules as they withdraw. These possibilities await further experimental dissection and analysis.

### 3. Conclusions and future perspectives

The simple observations and interpretations offered here provide a rationale for why such discontinuous or broken-apart PSDs could very likely represent the physical substrate of the temporary “tagging” of synapses that is generally thought to be so important for *initiating* the whole process of LTP (**Reference Set #19 (synaptic tagging in LTP).** This initiation is thought to prepare the synapse for later, long-term enhancement *via* new protein synthesis, which is presumed by all, to be the consolidating event that culminates LTP (**Reference Set #10 (Background of Long Term Potentiation), OP CIT).**

The rationale would be that the breakup of solid plaques or disks of PSD would create *new edges* where “modules” or molecular components of postsynaptic receptors, channels, and signaling molecules could be added, to enlarge or even create completely new PSD-plaques. In other words, that the discontinuities and irregularities in the PSD created by the act of perforation, due to the exacerbation of synaptic vesicle recycling in the presynapse, would create additional (and new) *free edges* around the plaques - - which originally had, by the simple fact that they were disk-like - - had the *minimum* amount of free edges possible, for a given collection of receptors.

The logical conclusion to draw from this, seems to us to be that newly-generated PSD “free edges” represent the synaptic “tags” that initiate LTP (**Reference Set #19 (synaptic tagging in LTP) OP CIT**). Although we did not attempt in this report to provide the complete structural evidence for this hypothesis, we predict that it should soon become available from many of the new LM- and EM-methods that are being developed to ‘tag’ various pre- and postsynaptic proteins and protein-complexes (**Reference Set #9 (synaptic ‘modules’ via LM-viewing) OP CIT**). Here, we focused only on providing direct EM-images that demonstrated how (and why) enhanced bursts of presynaptic secretory-activity apparently *create* or *cause* the perforation of otherwise plaque-like postsynaptic densities, in the first place.

## MATERIALS AND METHODS

### I. Cell-culture

Dissociated-cell hippocampal cultures were prepared from papain-dissociated hippocampi, which were harvested from embryonic day-20 rat fetuses, then plated onto confluent glial feeder-cultures on 22mm glass coverslips, and grown for 3-4 weeks before use. Throughout this growth-period, the culture-medium was half-exchanged 3x weekly with fresh medium containing MEM (with Earle’s salts, 6 g/l glucose, and 3.7 g/l sodium bicarbonate) supplemented with 5% (v/v) heat-inactivated horse serum, 2% (v/v) fetal bovine serum, and 2 mM Glutamax (all from Life Technologies), plus 136 uM uridine and 54 uM 2-deoxy-5-fluoro-uridine (from Sigma), plus N3-supplement from Sigma (which contains BSA, apotransferrin, putrescine, selenium, T3, insulin, progesterone, and corticosterone. (For further details, see Ransom et al., 1977, and Mayer et al., 1989, in **<u>Reference Set #11 (Methods for</u> <u>hippocampal slices & cultures</u>**). Throughout this time, the coverslips were maintained in P35 culture-dishes in a 36°C incubator with 10% CO2.

### II. Culture-treatments

To conduct the experiments, the culture-dishes containing the coverslips were removed from the CO2 incubator and *immediately* washed with a bicarbonate- and phosphate-free ‘Ringers’ solution… (removing bicarbonate so that the cultures did not alkalinize in room air, with its low CO2, and removing phosphate so that the subsequent application of lanthanum did not just precipitate as La(PO4)3). Henceforth, they were maintained on a rotating platform in a 37°C waterbath. After three more washes in HCO3- and PO4-free ‘Ringers,’ for a total time of 12min, the cultures were then exposed to a ‘Ringers’ solution containing 0.1mM LaCl3 (with its usual CaCl2 reduced from the usual 2mM CaCl2 to only 1mM, to minimize any competition of Ca++ with the La+++. Alternatively, cultures were treated for 5-15min at 37°C with high K+ (a ‘Ringers’ containing 90mM KCl (whose osmolarity had been compensated by reducing the concentration of NaCl)), or treated 5-10min at 37°C with 50–60μM of N-methyl-D-aspartic acid (NMDA) in normal HCO3-free ‘Ringers,’ but containing our usual 2mM of CaCl2 and 3mM of NaH2PO4.

### III. Fixation and processing

Primary fixation was accomplished by replacing the Ringers solution in the culture-dishes with 2% glutaraldehyde, freshly dissolved from a 50% stock (from EMS, Inc.) into a “substitute Ringer’s,” where the normal 5mM Hepes buffer concentration was increased 6x for fixation purposes (and NaCl was decreased commensurately, to keep the solution isotonic). Most important at this point was to wash away the La+++ and restore the normal 2mM calcium in the medium, to prevent any La+++ precipitates in the extracellular spaces of the cultures. (Indeed, this 2mM calcium is maintained *throughout* primary fixation and postfixation, because we believe it helps to minimize cellular membrane deterioration.)

Immediately after the exchange into the glutaraldehyde, the culture-dishes were placed on a vigorously rotating table, and the fixative was exchanged one or two more times, to ensure rapid and uniform fixation. (Even though this aldehyde fixation was probably complete in just a few minutes, we still left the cultures in fixative for another 1-2 hours, or even overnight, before initiating postfixation.)

The sequence of postfixation was as follows. (This was all done at room temperature, because we believe that cooling biological membranes to 4°C at any time during fixation damages cellular membranes.) First, the glutaraldehyde and Hepes buffer was washed away with 100mM cacodylate buffer, with two exchanges over a period of at least 15-30 minutes. (This buffer always contained the same 2mM Ca++, in this and all subsequent steps, so it will henceforth be termed “Cacodylate-Ca”.) Next, the cultures were postfixed with 0.25% OsO4 and 0.25% potassium ferocyanide in Cacodylate-Ca buffer (made up fresh, by mixing 0.5% OsO4 with 0.5% KFeCN6 immediately before use) for exactly 30 minutes, no longer. Then, after washing away the OsO4 with fresh Cacodylate-Ca buffer (for 5-10 min), the cultures were “mordanted” with 0.5% tannic acid (mw 1700) in Cacodylate-Ca buffer (making sure to use a batch of tannic acid from EMS or from Polysciences that did not precipitate over time), for 30 minutes only, no longer. Finally, after washing away the tannate with fresh Cacodylate-Ca buffer, the pH over the cultures was dropped by a brief wash in 100mM acetate buffer at pH 5.2, to prepare them for “block-staining” with 0.5% uranyl acetate in this acetate buffer (pH 5.2 being the natural pH of dissolved UA, anyway). Then, after this block-staining, they were very briefly washed again in acetate buffer to remove the UA, and finally progressively dehydrated with ethanol in the usual manner (sequential 5-10 min rinses in 50% 75%, 95%, and 100% ethanol).

### IV. Epoxy embedding and thin-sectioning

Thereafter, the coverslips were removed from the P35 culture-dishes to polypropylene bottles, where they could be embedded in Araldite 502 epoxy resin (the old “English Araldite”), via an intermediate transfer from ethanol into propylene oxide, then into 2/3rds Araldite+1/3 propylene oxide, etc. (The bottles were needed because the propylene oxide would have dissolved the original P35 culture dishes). Finally, the fully infiltrated cultures still in their polypropylene bottles were covered with a 10-12mm deep layer of freshly-prepared Araldite 502 epoxy resin and vacuum-embedded in a 70°C vacuum-oven, using a strong mechanical-pump to draw off all air until the Araldite formed small bubbles (from release of residual propylene oxide and ethanol), and after re-admission of air, were left for 24-48 hours to fully polymerize.

When fully hardened, the polypropylene bottles were removed the oven, broken with pliers to release the Araldite blocks, and the blocks were cut with a jeweler’s saw into pieces appropriate for mounting at the desired orientation the ultramicrotome. Finally, the original glass coverslips on which the cultures were grown were dissolved off of the Araldite by a brief (5-10) min dip into full-strength hydrofluoric acid (47%HF), followed by plenty of washes. Blocks were initially sectioned at 0.5-1.0 micron and stained with 1% toluidine blue and 1% sodium borate in water for 15 sec on a hotplate, to examine in the LM and to orient further block-trimming for thin-sectioning.

### V. Electron Microscopy

Thin sections were cut at 40nm to obtain the crispest views of membranes, at 90nm for best general overviews, and at 150-200nm to obtain 3-D information about the overall deployment of synapses in the cultures. Thin-sections were picked up on high-transmission fine-hexagonal 200-mesh copper grids (made in England by Guilder, LTD, and sold in the US by Ladd Industries, cat. no. G200HHC), after the grids had been coated with a silver-thin film of Formvar, and then carbon-coated with by 10 sec of vacuum-evaporated carbon, for maximum specimen stability. Finally, sections were stained for 5 min drops of 1% lead citrate (in a closed dish with NaOH pellets around to prevent CO2-precipitation of the lead).

They were then examined in a standard TEM operated at 80KV and mounted with the smallest available objective aperture, for maximum contrast (and maximum removal of chromatic aberration from the thicker sections). They were photographed with the highest resolution digital camera possible, regardless of sensitivity, since such Araldite sections were essentially indestructible and could tolerate endless electron-bombardment. (We generally used the AMT ‘BioSprint’ 29 Megapixel Camera, due to its many superior operating features, as well as its very clear 6.5k x 4.5k images.) The final digital images were processed and colorized with Photoshop, taking special advantage of its “high-pass” filter when very dark features happened to be located next to very light areas in the images, which made details hard to see. (We typically set the high-pass filter at 40 pixels for our 6500×4500 pixel AMT images, and layered this filtered image on top of the original image, at 50% density.)

## POSTSCRIPT

We find it absolutely marvelous that over forty-five years ago, Sally Tarrant and Aryeh Routtenberg, from Northwestern University’s Neuroscience Laboratory in Chicago, had the prescience and foresight to add the following *tiny and obscure footnote* to a fine paper of theirs, a paper in which they described and discussed ‘synaptic spinules’ in the brains of rats they had prepped for EM by perfusion with Karnovsky’s fixative**. That footnote said:

P.S. *”It is also possible that the ‘synaptic spinule’ represents an active synapse and that the presynaptic invagination represents the coalescence of synaptic vesicles and the coated vesicle a device for membrane recycling (Heuser and Reese, 1973). The spine apparatus might contribute to the postsynaptic membrane as it protrudes into the presynaptic membrane invagination.”*

**Tarrant SB & A. Routtenberg (1977)

The ‘synaptic spinule’ in the dendritic spine: EM study of the hippocampal dentate gyrus. Tissue & Cell. 9: 461-473. DOI: 10.1016/0040-8166(77)90006-4 PMID: 337572

## ACKNOWLEDGEMENTS

We especially thank Christine A. Winters (from the Laboratory of Neurobiology, NINDS, NIH) for kindly producing and supplying us with all the gorgeous and healthy hippocampal dissociated-cell cultures that were used in this study. We also thank Thomas S. Reese for providing work-space and reagents to carry out these experiments in his Laboratory of Neurobiology, NINDS, NIH. Finally, for our salary-support, we greatly thank Joshua Zimmerberg, head of the Section on Integrative Biophysics in the National Institute of Child Health and Human Development (NICHD).

## Authors’ contributions

This is a solo-author manuscript. The author wrote, read and approved the final manuscript.

## Availability of data and materials

The electron micrographs and other datasets generated and/or analyzed during the current study are available from the corresponding author upon request.

## Ethics approval and consent to participate

The animal protocol under which this work was carried out was approved by the Animal Use and Care Committee of the National Institute of Neurological Disorders and Stroke (NINDS) (Animal protocol Number: ASP1159), and conforms to all NIH guidelines.

